# Strengthening Medial Olivocochlear Feedback Reduces the Developmental Impact of Early Noise Exposure

**DOI:** 10.1101/2025.01.03.631257

**Authors:** Valeria C. Castagna, Luis E. Boero, Mariano N. Di Guilmi, Camila Catalano Di Meo, Jimena A. Ballestero, Paul A. Fuchs, Amanda M. Lauer, Ana Belén Elgoyhen, Maria Eugenia Gomez-Casati

**Affiliations:** Instituto de Investigaciones en Ingeniería Genética y Biología Molecular, Dr. Héctor N. Torres, Consejo Nacional de Investigaciones Científicas y Técnicas, 1428 Buenos Aires, Argentina; Instituto de Farmacología, Facultad de Medicina, Universidad de Buenos Aires, 1121 Buenos Aires, Argentina and Consejo Nacional de Investigaciones Científicas y Técnicas, Argentina; Universidad de Buenos Aires, Facultad de Ciencias Exactas y Naturales, Departamento de Fisiología y Biología Molecular y Celular, 1428 Buenos Aires, Argentina; Department of Otolaryngology-Head and Neck Surgery, Johns Hopkins University School of Medicine; Baltimore, Maryland 21205, United States; Department of Neuroscience, Johns Hopkins University School of Medicine, Baltimore, Maryland 21205, United States; Center for Functional Anatomy and Evolution, Johns Hopkins University, School of Medicine, Baltimore, Maryland 21205, United States

**Author notes:** **Corresponding Author:** María Eugenia Gómez-Casati, Ph.D., Instituto de Farmacología. Facultad de Medicina. Universidad de, Buenos Aires., Address: Paraguay 2155. C1121ABG, Buenos Aires, Argentina. **Conflict of Interest:** The authors declare no competing financial or non-financial interest. **Author Contributions:** V.C.C, L.E.B, M.N.D-G, P.A.F, A.M.L, A.B.E and M.E.G-C designed research; V.C.C, C. C.dM and J.A.B performed experiments; V.C.C analyzed data. M.E.G-C wrote the paper. All authors discussed the results, read, edited, and commented on the manuscript.

## Abstract

The early onset of peripheral deafness significantly alters the proper development of the auditory system. Likewise, exposure to loud noise during early development produces a similar disruptive effect. Before hearing onset in altricial mammals, cochlear inner hair cells exhibit spontaneous electrical activity that drives auditory circuit development. This activity is modulated by medial olivocochlear (MOC) efferent feedback through α9α10 nicotinic cholinergic receptors in inner hair cells. In adults, these receptors are restricted to outer hair cells, where they mediate MOC feedback to regulate cochlear amplification. Although the MOC system’s protective role to prevent noise-induced hearing loss in adulthood is well-established, its influence during early developmental stages-especially in response to exposure to loud noise-remains largely unexplored. In this study, we investigated the role of MOC feedback during early postnatal development using α9 knockout (KO) and α9 knock-in (KI) mice of either sex, which respectively lack or exhibit enhanced cholinergic activity. Our findings reveal that both increased and absent olivocochlear activity result in altered auditory sensitivity at the onset of hearing, along with long– range alterations in the number and morphology of ribbon synapses. Early noise exposure caused lasting auditory damage in both wild-type and α9KO mice, with deficits persisting into adulthood. In contrast, α9KI mice were protected from noise-induced damage, with no long-term effects on auditory function. These results highlight the increased susceptibility of the auditory system during early postnatal development. Moreover, they indicate that an enhanced MOC feedback shields the auditory system from noise damage during this period.

**SIGNIFICANCE STATEMENT:** Early development represents a sensitive window for shaping auditory function. We show that the medial olivocochlear system is critical for establishing normal ribbon synapse density and size; key features for proper hearing onset. We also show that the developing auditory system is especially vulnerable to loud noise, with early exposure causing more severe and lasting effects than similar noise later in life. Notably, enhancing α9α10 nAChR receptor activity during this early stage offers protection against noise-induced damage, revealing a time-sensitive opportunity to safeguard auditory development.

## Introduction

Cochlear and central auditory system development relies on tightly coordinated cellular and molecular events, with critical refinements during early postnatal life to optimize sound encoding (Sanes and Walsh, 1998). Noise exposure during this sensitive period can disrupt these mechanisms, damaging cochlear structures and neural circuits (Chang and Merzenich, 2003; Keuroghlian and Knudsen, 2007; Zhou and Merzenich, 2008; Sanes and Bao, 2009; Lauer et al., 2011). Early acoustic exposure affects neuronal sensitivity, innervation, frequency tuning and response characteristics across the auditory pathway (Caminos et al., 2005; Waguespack et al., 2007; Sanes and Bao, 2009; Dorrn et al., 2010; Sun et al., 2010; Froemke and Jones, 2011). Such disruptions can lead to permanent auditory deficits, emphasizing the importance of safeguarding the developing auditory system.

In altricial mammals, inner hair cells (IHCs) exhibit spontaneous electrical activity that propagates into the central nervous system (CNS) before hearing onset (∼P12 in mice) (Lippe, 1994; Marcotti et al., 2003; Sonntag et al., 2009; Tritsch and Bergles, 2010; Johnson et al., 2011; De Faveri et al., 2025). This activity is crucial for neuronal survival and for refining auditory circuits (Tierney et al., 1997; Friauf and Lohmann, 1999; Mostafapour et al., 2000; Leake et al., 2006; Clause et al., 2014; Di Guilmi et al., 2019). During this period, MOC neurons from the superior olivary complex form transient axosomatic synapses with immature IHCs, modulating their spontaneous firing (Glowatzki and Fuchs, 2000; Marcotti et al., 2004; Johnson et al., 2013). These synapses, mediated by α9α10 nicotinic cholinergic receptors (nAChRs) coupled to SK2 channels, hyperpolarize IHCs (Glowatzki and Fuchs, 2000; Marcotti et al., 2004; Goutman et al., 2005), shaping burst patterns that guide synaptic development and ensure bilateral representation in the CNS (Walsh et al., 1998; Johnson et al., 2013; Clause et al., 2014; Di Guilmi et al., 2019; Wang et al., 2021). As IHCs mature, they lose direct efferent contacts and no longer respond to acetylcholine (ACh) (Katz et al., 2004; Roux et al., 2011). At this stage, MOC fibers cross the tunnel of Corti and innervate OHCs to regulate cochlear amplification (Wheeler et al., 1994; Ernfors et al., 1995; Knipper et al., 1995; Simmons et al., 1996; Walsh et al., 1998; Vattino et al., 2020). ACh-induced responses in OHCs emerge by P6, shifting from nicotinic-like to outward potassium currents by hearing onset (Dulon and Lenoir, 1996; He and Dallos, 1999). The olivocochlear system matures rapidly within the first 2–3 postnatal weeks, with adult-like innervation by the end of the first month (Pujol et al., 1978; Pujol, 1985; Simmons et al., 1990, 1996). While the protective role of the MOC system in preventing noise-induced hearing loss in adulthood is well-documented (Liberman, 1991; Maison and Liberman, 2000; Maison et al., 2002, 2013; Taranda et al., 2009; Boero et al., 2018), its impact during early developmental stages, remains largely unexplored.

This study explores how cholinergic MOC input strength influences auditory development and noise vulnerability during this sensitive period. We used mice with altered α9α10 nAChR activity: α9KO mice, which lacks efferent transmission (Vetter et al., 1999), and α9KI, carrying an α9 point mutation that leads to an enhanced MOC effect (Taranda et al., 2009). We show that MOC strength modulates hearing onset timing and IHC ribbon synapse formation. Wild-type (WT) mice exposed to noise at hearing onset showed greater threshold shifts and synaptic loss than similar exposure in adulthood (Boero et al., 2018), revealing heightened developmental vulnerability. In α9KO mice, early exposure resulted in a persistent loss of auditory nerve synapses, emphasizing the essential role of the MOC pathway. Conversely, α9KI mice were protected from both thresholds shifts and synaptic loss after early noise exposure. This protection occurred during a developmental window when the auditory system is still maturing and MOC-OHC synapses are forming (Simmons et al., 1996; Vattino et al., 2020). These findings highlight the importance of efferent cholinergic input during cochlear development and suggest that strengthening MOC feedback during this window can mitigate long-term effects of early acoustic trauma.

## Materials and Methods

### Animals

α9KO and α9KI mice have been previously described (Vetter et al., 1999; Taranda et al., 2009) and were backcrossed with the congenic FVB.129P2-*Pde6bþ Tyrc-ch/AntJ* strain (https://www.jax.org/strain/004828) for seventeen generations (i.e., N-17). We maintained a consistent male-to-female ratio across all experimental groups and genotypes. No sex-related differences were observed in the data. This study included adult and pup animals aged P12 to P75, which were housed in a sound-controlled room within the animal care facility. Background sound levels in this environment were consistently below 50 dB SPL. Noise levels were measured using an electret condenser microphone (FG-23329-PO7; Knowles). Transient sounds during routine cage cleaning and daily maintenance by animal care staff were minimal, with peaks not exceeding 70 dB SPL. The methods reported in this study adhere to the ARRIVE guidelines for reporting *in vivo* animal research (https://arriveguidelines.org/), ensuring transparency and reproducibility. All procedures were conducted in accordance with institutional and ethical guidelines for animal care and use.

### Auditory function tests

Physiological assessments of cochlear and central function, including auditory brainstem responses (ABRs) and distortion-product otoacoustic emissions (DPOAEs), were performed in mice anesthetized with xylazine (10 mg/kg, i.p.) and ketamine (100 mg/kg, i.p.) and placed in a soundproof chamber maintained at 32°C to counteract the drop in body temperature caused by anesthesia. ABR represents sound-evoked potentials generated by neural circuits in the ascending auditory pathways, while DPOAEs are sounds emitted by the cochlea, specifically by OHCs, in response to two pure-tone stimuli presented simultaneously. Sound stimuli were delivered through a custom acoustic system with two dynamic earphones used as sound sources (CDMG15008–03A; CUI) and an electret condenser microphone (FG-23329-PO7; Knowles) coupled to a probe tube to measure sound pressure near the eardrum. Data collection was conducted using a National Instruments PXI-based system equipped with 24-bit input/output boards and operated through a custom LabView software interface. For ABRs, platinum needle electrodes were placed into the skin at the dorsal midline close to the neural crest and pinna with a ground electrode near the tail. ABR potentials were evoked with 5 ms tone pips (0.5 ms rise-fall, with a cos^2^ envelope, at 40/s) delivered to the eardrum at log-spaced frequencies from 5.6 to 45.25 kHz. The response was amplified 10,000X with a 0.3–3 kHz passband. Sound level was raised in 5 dB steps from 20 to 80 dB sound pressure level (SPL). At each level, 1024 responses were averaged with alternated stimulus polarity. Threshold for ABR was defined as the lowest stimulus level (<80 dB SPL) at which a repeatable wave 1 could be identified in the response waveform. The ABR wave 1 amplitude was computed by off-line analysis of the peak-to-peak amplitude of stored waveforms. The DPOAEs in response to two primary tones of frequency f1 and f2 were recorded at 2f1-f2, with f2/f1=1.2, and the f2 level 10 dB lower than the f1 level. Ear-canal sound pressure was amplified and digitally sampled at 4 µs intervals. DPOAE threshold was defined as the lowest f2 level in which the signal to noise floor ratio is >1.

### Noise exposure

Animals were exposed under anesthesia at P15 to broadband noise (1–16 kHz, 100 dB SPL) for 1 hour using closed-field earphones (CDMG15008–03A; CUI) as the sound source. This broader frequency range was selected to induce significant threshold shifts in the apical region of the cochlea, as previously described in (Boero et al., 2018). Noise exposure was conducted in the same acoustic chamber used for cochlear function tests. The noise consisted of a flat-spectrum broadband signal calibrated to the target SPL immediately before each exposure session.

### Animal monitoring and health assessment

To assess the impact of repeated anesthesia on the health and well-being of the animals, their weight and motor activity were monitored daily throughout the auditory function recordings and noise exposure protocols. Animals were observed during their recovery from anesthesia to ensure no adverse effects on their general condition. Growth curves showed no significant differences in weight between animals that were anesthetized and untreated controls animals, suggesting that anesthesia did not affect normal growth. Furthermore, no behavioral changes or motor impairments were observed in the anesthetized animals when compared to controls, indicating that repeated anesthesia had no detectable impact on overall health or behavior.

### Cochlear processing and immunostaining

Cochleae were perfused intralabyrinthly with 4% paraformaldehyde (PFA) in phosphate-buffered saline (PBS), post-fixed with 4% PFA overnight, and decalcified in 0.12 M EDTA. Cochlear tissues were microdissected and permeabilized by freeze/thaw cycles in 30% sucrose. The dissected samples were then separated into distinct wells based on cochlear region: apex, middle and base. Tissue samples were blocked with 5% normal goat serum and 1% Triton X-100 in PBS for 1 h, followed by incubation in primary antibodies (diluted in blocking buffer) at 37°C for 16 h. The primary antibodies used in this study were: 1) anti-C-terminal binding protein 2 (mouse anti-CtBP2 IgG1; BD Biosciences, San Jose, CA; Cat#612044, RRID:AB_399431, 1:200) to label the presynaptic ribbon, 2) anti-glutamate receptor 2 (mouse anti-GluA2 IgG2a; Millipore, Billerica, MA; Cat#MAB397, RRID:AB_11212990, 1:2000) to label the post-synaptic receptor plaques and 3) anti-Myosin 7a (rabbit anti-MyosinVIIa, Proteus Biosciences; Cat# 25-6790, RRID:AB_10015251, 1:200) to label the cochlear hair cells. Tissues were then incubated with the appropriate Alexa Fluor-conjugated fluorescent secondary antibodies (Invitrogen, Carlsbad, CA; 1:500 in blocking buffer) for 2 hours at room temperature. Finally, tissues were mounted on microscope slides in FluorSave mounting media (Millipore, Billerica, MA).

### Confocal microscopy and image processing

For IHC synaptic counts, cochlear tissue from the apex, middle and base regions was first imaged at low magnification to correlate each piece with its corresponding frequency region (Müller et al., 2005). The apical region corresponds to 4– 12 kHz, the middle region to 12–20 kHz, and the base region to 22–40 kHz. Z-stacks images were then acquired using a Leica TCS SPE microscope with a 63X oil-immersion lens (1.5X or 4X digital zoom). Imaging settings were consistent across all samples, ensuring no pixel saturation. For each stack, the Z-step was set to 0.3 µm, with a pixel size of 0.11 µm in both the X and Y axes. Each stack typically contained 10 to 20 IHCs. Image stacks were processed using Fiji software (RRID:SCR_002285) (Schindelin et al., 2012), and a custom plugin was developed to automate the counting of synaptic ribbons, glutamate receptor patches and co-localized synaptic puncta. The algorithm’s accuracy was validated by comparing automated counts to manual counts, ensuring reliable quantification. Briefly, each channel was analyzed separately, and maximum projections were generated to quantify the number of CtBP2 or GluA2 puncta. Additionally, a composite between the three channels was produced to draw the different regions of interest (ROI) that correspond to each IHC taking the myosin staining as a reference. The maximum projections from the single channels were multiplied to generate a merged 32-bit image. Then, they were converted to binary images after a custom thresholding procedure. An automatic counting of the number of particles on each ROI was performed. The plugin used for synapse quantification, along with detailed instructions, is available at: https://github.com/vcastagna/CountsSynapses.

For quantitative volumetric synaptic analysis, different regions from the whole cochlea were imaged and confocal z-stacks were processed in Fiji. z-stacks were deconvolved applying 10 iterations of the Richardson-Lucy algorithm in the DeconvolutionLab2 plugin (Sage et al., 2017), an algorithm that has been used previously in cochlear whole mounts (Jing et al., 2013).

Then, the 3D Object Counter plugin was used to perform three-dimensional image segmentation and object individualization of the deconvoluted stacks. According to previous work (Liberman et al., 2011; Gilels et al., 2013; Paquette et al., 2016) we reasoned that a synapse would occupy a volume between 0.04 µm3 to 5 µm3, and then we set a size range of 20-2000 voxels, with a voxel size of 0.11 µm x 0.11 µm x 0.2 µm. The volumes of presynaptic ribbons (CtBP2 fluorescence) and postsynaptic AMPA receptor clusters (GluA2 fluorescence) at IHC-afferent synapses were measured using the 3D ROI Manager (Ollion et al., 2013). All exported volumes were checked to ensure that volumes were within instrument resolution limits after threshold adjustments.

### Statistical analysis

Data was analyzed using R Statistical Software (RRID:SCR_001905). Shapiro-Wilks test was used for testing normal distribution of the residuals. In cases where the data did not follow a normal distribution, the following tests were used: Kruskal-Wallis test followed by multiple contrasts according to Dunn’s method for comparisons between three groups (IHCs synaptic counts in the control condition, ABR and DPOAE thresholds during development) and Mann-Whitney U test for comparisons between two groups (ABR thresholds, DPOAE thresholds, and IHCs synaptic counts). If data were normally distributed, a one-way ANOVA was conducted, followed by Holm-Sidak post hoc tests for three-group comparisons (wave 1 amplitudes and synaptic volumes in the control condition). For two-group comparisons, a t-test was used (synaptic volumes). Statistical significance was set to p<0.05.

## Results

### Onset of auditory perception in mice with varying levels of efferent inhibition

In this study we used mice with distinct levels of α9α10 nAChR activity: 1) WT, with normal MOC function; 2) α9KO, which lacks cholinergic transmission between MOC neurons and hair cells (Vetter et al., 1999), and 3) α9KI mice carrying an α9 nAChR subunit point mutation that leads to enhanced responses to ACh (Taranda et al., 2009). To explore the impact of noise exposure at hearing onset, we first identified the optimal developmental stage for introducing such exposure, by assessing the onset of hearing under control (unexposed) conditions in these mouse models. This was done by tracking auditory brainstem responses (ABRs), the sound-evoked potentials generated by neuronal circuits in the ascending auditory pathway, at different postnatal ages across the three groups. Individual ABR waves reflect the activation of auditory periphery and brainstem processing relay stations within the first ∼7 ms after sound stimulus onset (Buchwald and Huang, 1975; Karplus et al., 1988; Shapiro, 1988; Shaw, 1988; Melcher and Kiang, 1996; Kim et al., 2013). Wave 1 represents the synchrony in firing of spiral ganglion neurons and is highly correlated with the number of synapses between IHCs and auditory nerve fibers (Buchwald and Huang, 1975; Antoli-Candela and Kiang, 1978; Kujawa and Liberman, 2009). Measurements were taken daily from P12 to P16, as well as P22 and P75, in anesthetized mice. Interestingly, we found that ABR onset occurred earlier in α9KI mice, with enhanced MOC function, compared to WT mice with normal MOC synapses and was delayed in α9KO mice lacking α9α10 nAChR activity (Figure 1a and b). Representative ABR traces recorded at P14 in response to a 16 kHz stimulus across varying intensities illustrate differences in ABR thresholds among the three mouse groups with distinct levels of α9α10 nAChR activity (Figure 1a). At this developmental stage, α9KI mice exhibited lower ABR thresholds, and more distinct peaks at suprathreshold intensities compared to WT mice. In contrast, α9KO mice showed higher ABR thresholds relative to WT mice (Figure 1a). Figure 1b shows the mean ABR thresholds measured at test frequencies of 11.33, 16, and 22.65 kHz from P12 to P75 across the three groups of mice. A comparison between α9KI and WT mice revealed that α9KI mice displayed measurable ABR responses as early as P13 (Kruskal-Wallis, followed by Dunn’s test, degree of freedom [df] = 2, p = 0.01 at 22.65 kHz; Figure 1b) and achieved mature threshold levels earlier than their WT littermates (Kruskal-Wallis, followed by Dunn’s test, df = 2, p = 0.02 at 16 and 22.65 kHz at P14; Figure 1b). In contrast, ABR onset was delayed by a day in α9KO compared to WT mice (Kruskal-Wallis followed by Dunn’s test, df = 2, p = 0.04 at 11.33 and 16 kHz; Figure 1b), indicating that cochlear maturation is slower in the absence of pre-hearing efferent modulation. To summarize, at P13, no ABR responses were observed in WT (0 out of 9) or α9KO (0 out of 7) mice in response to tone pips at 80 dB SPL across 11.33, 16, and 22.65 kHz. In contrast, 6 out of 9 α9KI mice exhibited ABR responses at P13. By P14, ABR responses were present in 77.7% of WT mice at 11.33 and 22.65 kHz and in 100% at 16 kHz. Meanwhile, 85% of α9KO mice responded at all frequencies, but with higher thresholds than WT (Figure 1b). All α9KI mice exhibited robust responses at all tested frequencies by P14. By P15, 100% of WT and 91% of α9KO mice showed ABR responses.

**Figure 1:**
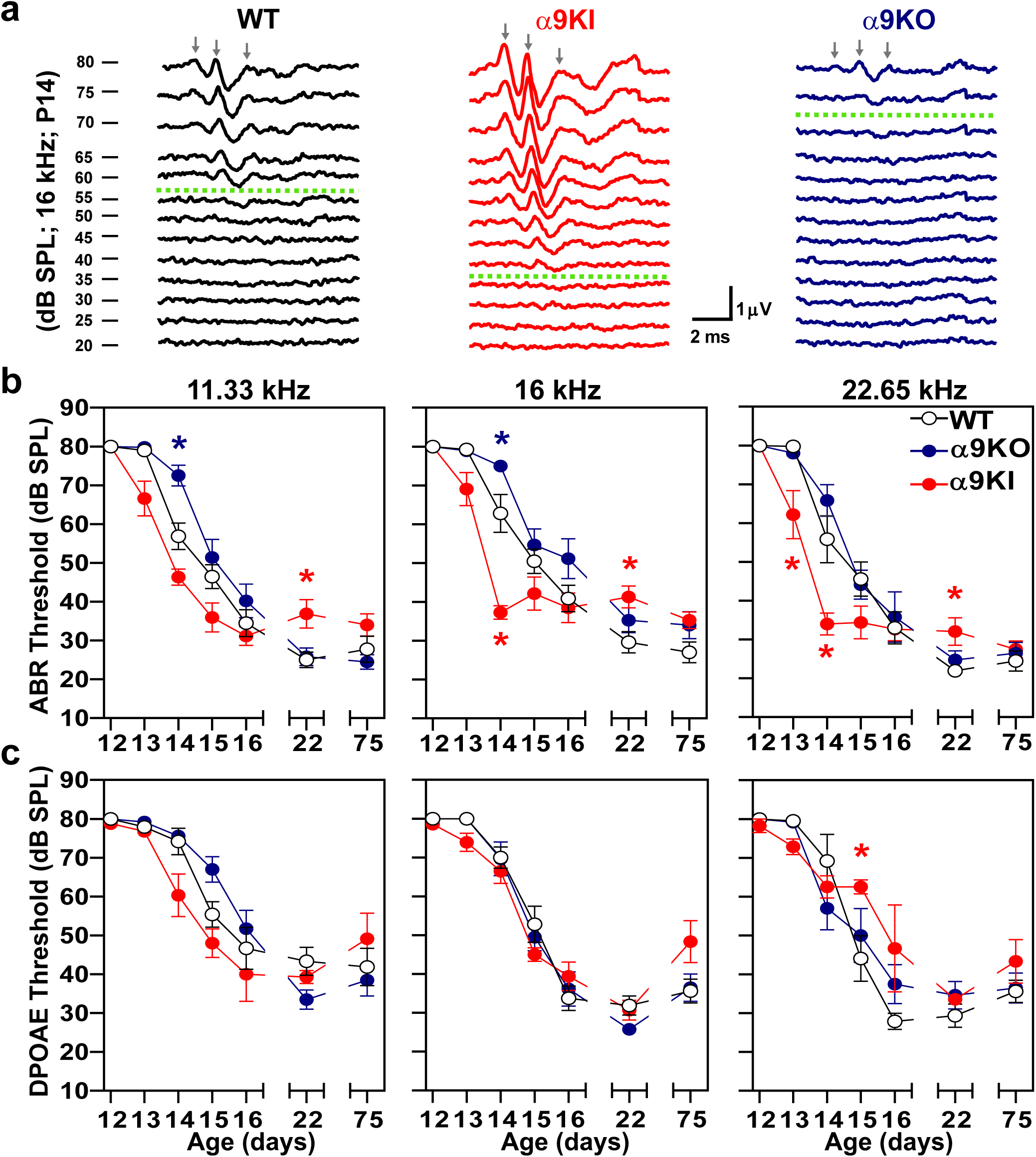
Auditory function in WT, α9KO, and α9KI mice across different postnatal ages. (**a**) Representative ABR waveforms from WT (black), α9KI (red), and α9KO (blue) mice recorded at P14 in response to 16 kHz tone bursts at increasing sound pressure levels (dB SPL). Thresholds, defined by the presence of wave 1, are marked by green dashed traces. Waves 1, 2 and 3 are indicated by gray arrows. The scale bar applies to all three recordings. (**b**) Mean ABR thresholds from P12 to P75 in WT (P12 to P14 n = 9 and P15 to P75 n = 14), α9KO (P12 to P14 n = 7 and P15 to P75 n = 11), and α9KI (P12 to P14 n = 9 and P15 to P75 n = 12) mice at 11.33, 16, and 22.65 kHz. ABR waveforms first became detectable at P13 in α9KI mice, at P14 in WT mice, and between P14 and P15 in α9KO mice. (**c**) Mean DPOAE thresholds from P12 to P75 in WT (P12 to P14 n = 9 and P15 to P75 n = 14), α9KO (P12 to P14 n = 7 and P15 to P75 n = 11) and α9KI (P12 to P14 n = 9 and P15 to P75 n = 12) mice at 11.33, 16 and 22.65 kHz. DPOAE thresholds showed the same trend as ABR only at 11.33 kHz. Group means ± SEM are shown. Red asterisks represent the statistical significance of α9KI compared with WT, and blue asterisks represent the statistical significance of α9KO compared with WT (Kruskal-Wallis followed by post hoc Dunn test, *p < 0.05).

Tone pip-evoked ABR wave 1 amplitudes were then analyzed at a suprathreshold level of 80 dB SPL at the different test frequencies in the three groups of mice with varying levels of α9α10 nAChR activity (Figure 2a, b). As illustrated in the representative traces in Figure 2a, measurable ABR wave 1 amplitudes at P13 were only observed in α9KI mice. At P14, the number of discernible peaks were fewer in WT and α9KO mice compared to α9KI mice. Up to P15, response peaks at 80 dB SPL were slightly larger and more distinct in α9KI mice, reflecting enhanced auditory responses during this postnatal developmental window. However, starting at P16, α9KI mice exhibited smaller ABR wave 1 amplitudes, with significant differences observed at P75 at 16 and 22.65 kHz compared to WT mice (one way ANOVA followed by Holm-Sidak test, df = 2, p = 0.03 at both frequencies; Figure 2b). In α9KO mice, wave 1 amplitudes remained smaller than those of WT and α9KI mice until P16, with the difference being significant at 22.65 kHz (one way ANOVA followed by Holm-Sidak test, df = 2, p = 0.02; Figure 2b). These findings align with the ABR threshold results, indicating that the strength of efferent cholinergic inhibition to hair cells contributes to the timely development of auditory function. The reduced ABR wave 1 amplitudes obtained in mature mice with either enhanced or null cholinergic activity indicates that proper MOC modulation during development might be crucial for the correct formation of auditory nerve synapses. Analysis of ABR wave 1 latencies at different postnatal ages showed no significant differences between genotypes (Figure 2c). While wave 1 latencies were consistently longer across all groups at P14, they shortened to levels typical of mature mice by P15.

**Figure 2:**
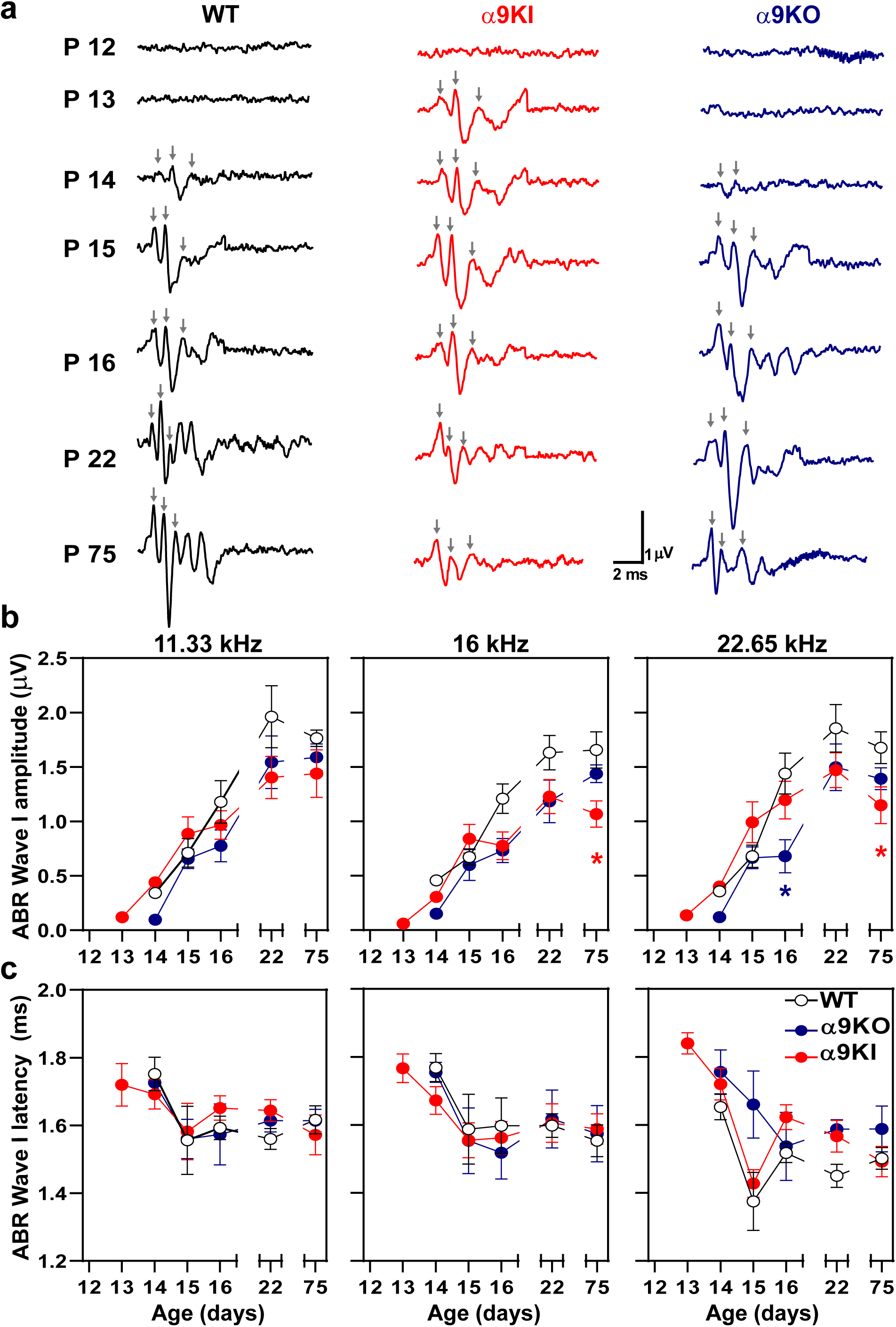
Suprathreshold ABR Wave1 amplitudes and latencies in WT, α9KO, and α9KI mice during postnatal development. (**a**) Representative examples of ABR waveforms recorded at 16 kHz, 80 dB SPL, from WT (black), α9KI (red), and α9KO (blue) mice. Waves 1, 2 and 3 are indicated by gray arrows. Each column illustrates waveform maturation across developmental stages (P12–P16, P22, P75) for each genotype. At P13, α9KI mice exhibit measurable responses, while distinct ABR waves emerge in WT and α9KO mice by P14. Waveform morphology stabilizes by P22 and remains consistent through P75. Scale bars apply to all traces. (**b**) ABR wave 1 amplitudes (µV) at 80 dB SPL for 11.33, 16, and 22.65 kHz across developmental ages in WT, α9KO, and α9KI mice. Amplitudes were measured as the difference between the maximum and minimum peaks. No values are shown for P12 due to the absence of discernible waves. (**c**) ABR wave 1 latencies (ms) at the same frequencies. A developmental decrease in latency is observed between P14 and P15 in all genotypes, though not statistically significant. The number of animals per genotype is the same as in Figure 1. Data represent mean ± SEM. Red asterisks denote significant differences between α9KI and WT; blue asterisks indicate significant differences between α9KO and WT (one-way ANOVA followed by Holm-Sidak test, *p < 0.05; **p < 0.01).

OHC function was assessed through distortion-product otoacoustic emissions (DPOAE) at the different postnatal ages. In a healthy cochlea, two close pure tones produce distortions due to OHC nonlinearities, which are amplified by OHC electromotility and detected with a microphone in the external ear canal (Shera and Guinan Jr., 1999; Robles and Ruggero, 2001). DPOAE responses followed a similar pattern to ABR thresholds, though the differences were not statistically significant (Kruskal-Wallis, df = 2, p > 0.05 at all frequencies; Figure 1c). At P13, DPOAE responses were nearly absent in WT (1 out of 9) and α9KO (0 out of 7) mice across the tested frequencies. In contrast, 3 out of 9 α9KI mice showed DPOAE responses at 16 and 22.65 kHz. By P14, DPOAE responses were detectable in 75% of WT and α9KO mice at 16 and 22.65 kHz, and in 60% of mice at 11.33 kHz. All α9KI mice exhibited strong DPOAE responses at all tested frequencies by P14. By P15, response rates reached 100% in both WT and α9KO mice. Note that at P22 and P75, mean ABR and DPOAE thresholds in α9KI mice were slightly elevated by 5–10 dB compared to WT and α9KO mice (Figure 1b, c). This elevation reached statistical significance only for ABR thresholds at P22 (Kruskal-Wallis, df = 2, p = 0.03 at 11.33 kHz; p = 0.04 at 16 kHz and p = 0.02 at 22.65 kHz). The elevated thresholds in α9KI mice at these mature stages are most likely due to enhanced suppression of OHCs activity resulting from increased cholinergic neurotransmission. This effect, previously described by Taranda et al. (2009), can be reversed by strychnine, a known antagonist of the α9α10 nAChR. Collectively, these findings indicate that the onset of hearing is altered in mice with either absent or enhanced olivocochlear cholinergic feedback.

### ABR thresholds after noise exposure at hearing onset in mice with different degree of MOC inhibition

To investigate the impact of noise exposure during early development, we exposed WT, α9KO, and α9KI mice to loud noise at P15. This allowed us to evaluate how acoustic experience influences auditory function before full threshold maturation in each genotype. Mice were exposed to 1–16 kHz noise at 100 dB SPL for 1 hour at P15. Auditory responses were then assessed at 1-, 7-, and 60-days post-exposure to track functional changes over time.

Previous studies in WT mice exposed to the same noise protocol at P21 showed a significant increase in auditory thresholds of 10–35 dB SPL one day after exposure, which returned to baseline within a week, indicating a transient threshold shift (Boero et al., 2018). In contrast, as shown in Figure 3a, no changes in ABR thresholds were observed at P16, one day after noise exposure, across frequencies of 8, 16, and 22.65 kHz in WT mice compared to unexposed controls. Remarkably, ABR thresholds at P16 were slightly lower than those measured at P15, prior to noise exposure, and showed no significant difference when compared to unexposed controls at the same age. This suggests that noise did not have an immediate effect on the maturation of auditory thresholds at this postnatal developmental stage. One week after exposure, however, ABR thresholds were elevated at all frequencies in noise-exposed WT animals, and this increase persisted at the same level for at least two months post-exposure (Mann-Whitney test, df = 1, P22: p = 0.006 at 8 kHz; p = 0.007 at 16 kHz; p = 0.002 at 22.65 kHz and P75: p = 0.04 at 8 kHz; p = 0.03 at 16 kHz and p = 0.04 at 22.65 kHz; Figure 3a). This outcome contrasts with the transient threshold elevation observed in WT mice from the same genetic background, exposed at P21 using the same noise protocol (Boero et al., 2018), suggesting that noise exposure at this early developmental stage may disrupt the normal maturation process, potentially leading to more profound and lasting auditory deficits.

**Figure 3:**
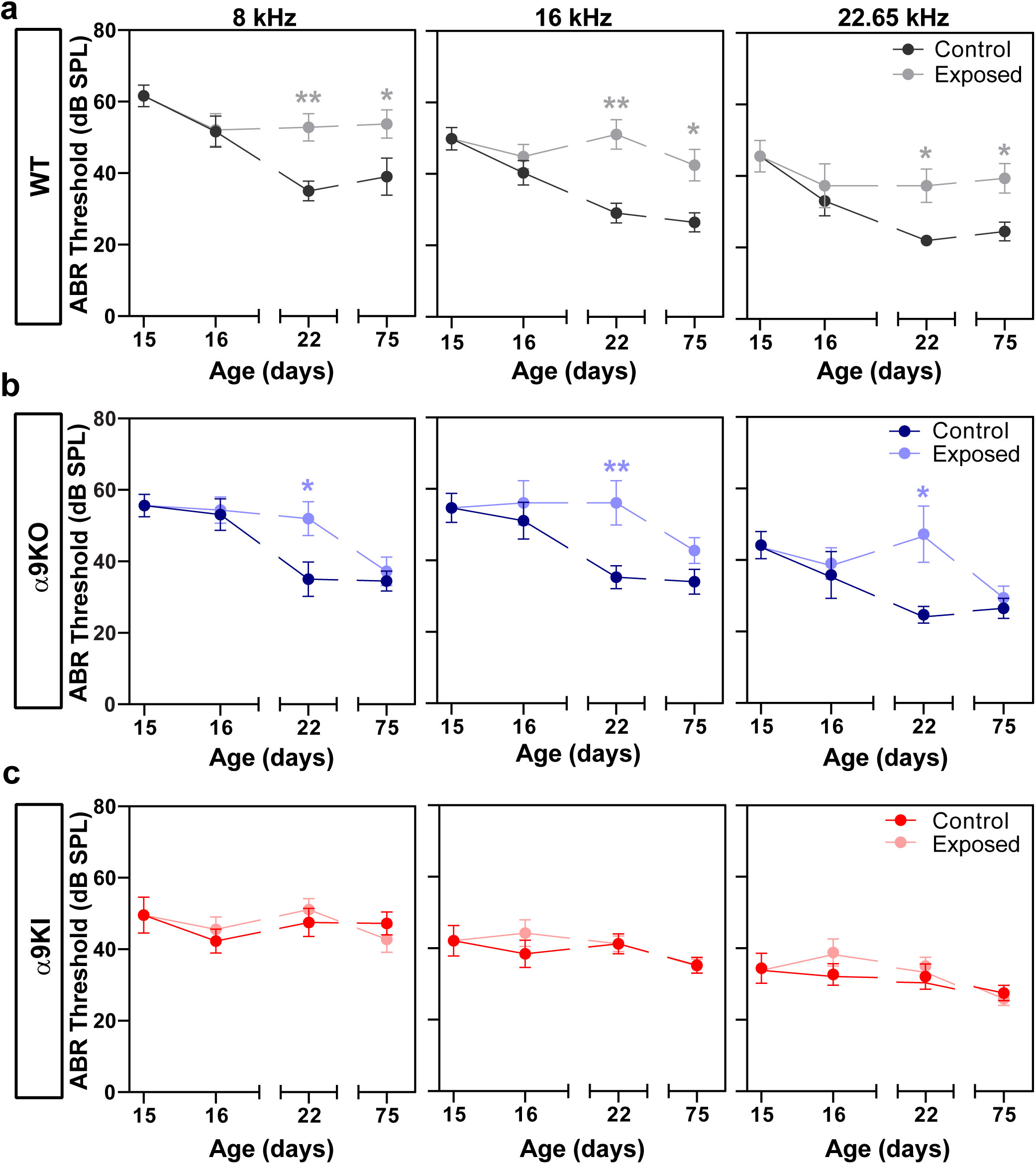
ABR wave 1 threshold measurements in control and noise-exposed WT, α9KO, and α9KI mice at different postnatal ages. ABR thresholds before and after exposure to noise at P15 in WT (n_control_ = 14 and n_Exposed_ = 12) (**a**), α9KO (n_control_ = 11 and n_Exposed_ = 11) (**b**), and α9KI (n_control_ = 12 and n_Exposed_ = 12) (**c**) mice at 8, 16 and 22.65 kHz. WT mice showed a significant increase in ABR thresholds 1 week and 2 months after exposure to noise, indicating a permanent threshold elevation. A recovery of ABR thresholds was observed in α9KO mice 2 months after exposure to noise. α9KI mice did not present any changes in ABR thresholds at any time after acoustic trauma. Each point represents the mean ± SEM. Asterisks represent the statistical significance (Mann-Whitney test, *p < 0.05; **p < 0.01).

A similar trend as seen in WT mice was observed after exposing α9KO mice to noise at P15 (Figure 3b). One day after exposure, there were no changes in ABR thresholds at 8, 16 and 22.65 kHz. However, seven days post-exposure, there was an elevation in auditory thresholds across all frequencies (Mann-Whitney test, df = 1, p = 0.03 at 8 kHz; p = 0.01 at 16 kHz and p = 0.02 at 22.65 kHz), which, unlike in WT mice, returned to normal levels after two months (Mann-Whitney test, df = 1, p > 0.05 at all frequencies; Figure 3b). Notably, in α9KI mice, ABR thresholds were not affected by noise exposure at all the frequencies tested (Figure 3c), suggesting that at this early age, the enhancement of cholinergic activity on OHCs provides protection from noise-induced cochlear threshold elevations (Mann-Whitney test, df = 1, p > 0.05 at all frequencies).

### Amplitudes of cochlear sound-evoked potentials after noise exposure at hearing onset in mice with different degree of efferent inhibition

To assess the neural activity across auditory nerve fibers, we measured ABR wave 1 amplitudes at frequencies of 8, 16, and 22.65 kHz at different ages after noise exposure in the three mouse groups (Figure 4). One day after noise exposure, we observed a reduction in wave 1 amplitudes across all frequencies in WT mice. Further testing of WT mice one week and two months post-exposure revealed a significant and long-lasting reduction, indicating a permanent decrease of ABR wave 1 amplitude (Mann-Whitney test, df = 1, P22: p = 0.007 at 8 kHz; p = 0.005 at 16 kHz; p = 0.005 at 22.65 kHz and P75: p = 0.04 at 8 kHz; p = 0.002 at 16 kHz and p = 0.002 at 22.65 kHz; Figure 4a). Similar permanent reductions in wave 1 amplitude were observed in α9KO mice after exposure to noise at 16 and 22.65 kHz (Mann-Whitney test, df = 1, P22: p = 0.04 at 16 kHz; p = 0.001 at 22.65 kHz and P75: p = 0.03 at 16 kHz and p = 0.04 at 22.65 kHz; Figure 4b). Notably, no changes in ABR wave 1 amplitudes were observed at any age post-exposure in mice with enhanced α9α10 nAChR activity, suggesting a preservation of cochlear synapses after noise exposure (Mann-Whitney test, df = 1, p > 0.05 at all frequencies; Figure 4c).

**Figure 4:**
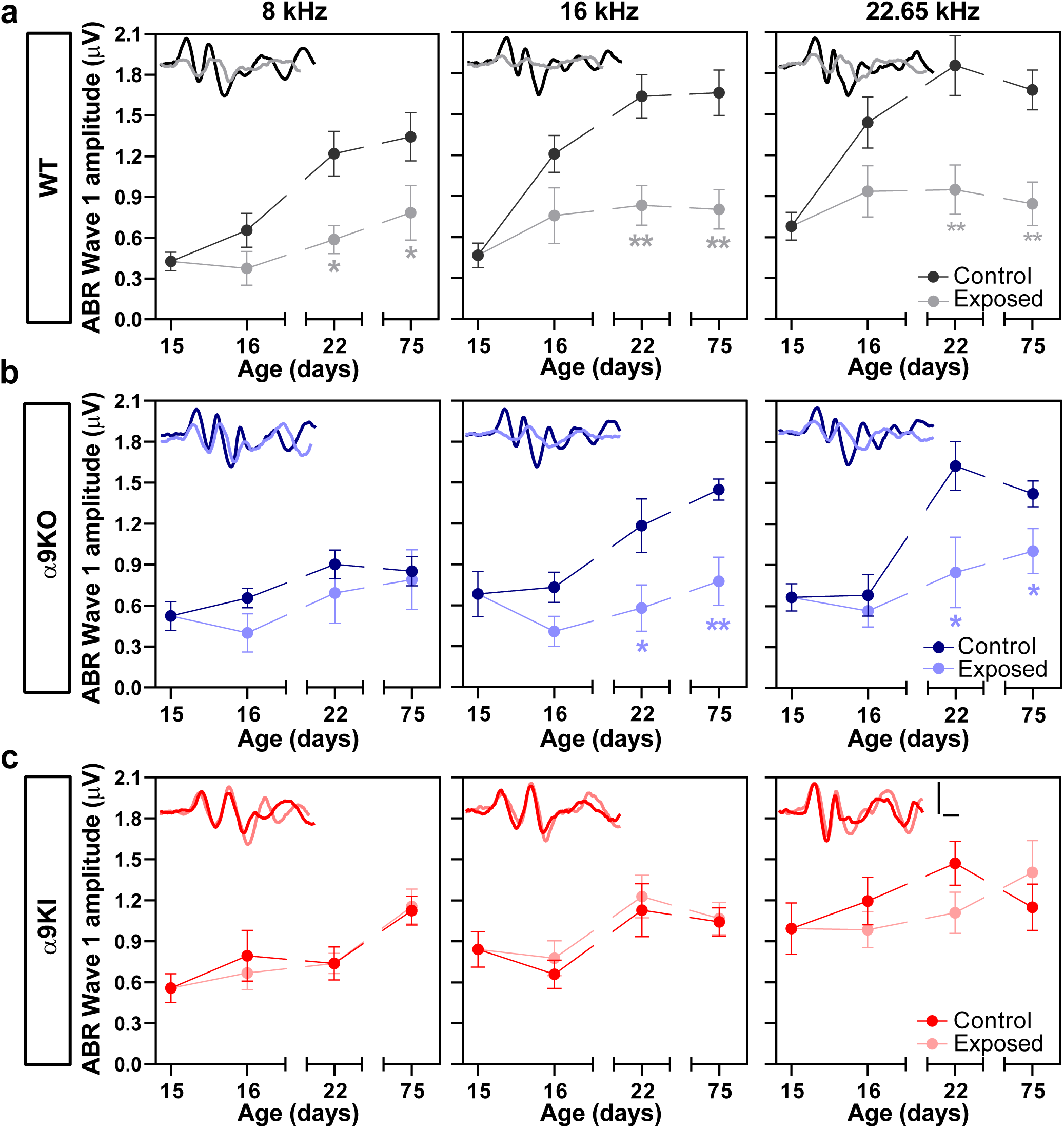
Suprathreshold response amplitude for ABR wave 1 after P15 noise exposure in WT, α9KO, and α9KI mice at different postnatal ages compared to unexposed controls. ABR wave 1 amplitudes (in µV) at 80 dB SPL at 8, 16 and 22.65 kHz in WT (n_Control_ = 14 and n_Exposed_ = 12) (**a**), α9KO (n_Control_ = 11 and n_Exposed_ = 11) (**b**), and α9KI (n_Control_ = 12 and n_Exposed_ = 12) (**c**) mice at the same time points shown in Figure 3. Each panel inset displays representative ABR waveforms recorded at 80 dB SPL for the corresponding frequency, shown for all groups at P75. Scale bars (1 μV vertical, 1 ms horizontal) apply to all traces. WT and α9KO mice exhibited reduced ABR wave 1 amplitudes following noise exposure, whereas wave 1 amplitudes in α9KI mice remained unaffected. Data are presented as mean ± SEM. Statistical significance is indicated by asterisks (Mann-Whitney test, *p < 0.05; **p < 0.01).

### DPOAE thresholds after noise exposure at hearing onset in mice with different levels of α9α10 nAChR activity

To assess the impact of hearing onset noise exposure on the function of OHCs, we recorded DPOAEs at 8, 16, and 22.65 kHz on the three groups of mice with varying levels of MOC function (Figure 5). During early development, the auditory system undergoes significant maturation, reflected by a progressive decrease in DPOAE thresholds with age, as shown in Figure 1c. In WT mice, one day after noise exposure at P16, there was no difference in DPOAE thresholds between the noise-exposed group and unexposed controls (Figure 5a). However, when comparing DPOAE thresholds at P16 to those at P15 within both groups (i.e. exposed and unexposed controls), a clear reduction was observed (Figure 5a). This decline in thresholds underscores the natural maturation process of the auditory system, indicating that early noise exposure does not immediately disrupt the ongoing development of DPOAE thresholds at this early postnatal stage. By P22, seven days after noise exposure, an increase in DPOAE thresholds was observed that was statistically significant only at 16 kHz (Mann-Whitney, df = 1, p = 0.01; Figure 5a). Yet, two months later, DPOAE thresholds were comparable to those of unexposed WT controls across all tested frequencies, suggesting that early noise exposure did not result in long-term changes in OHC function (Figure 5a).

**Figure 5:**
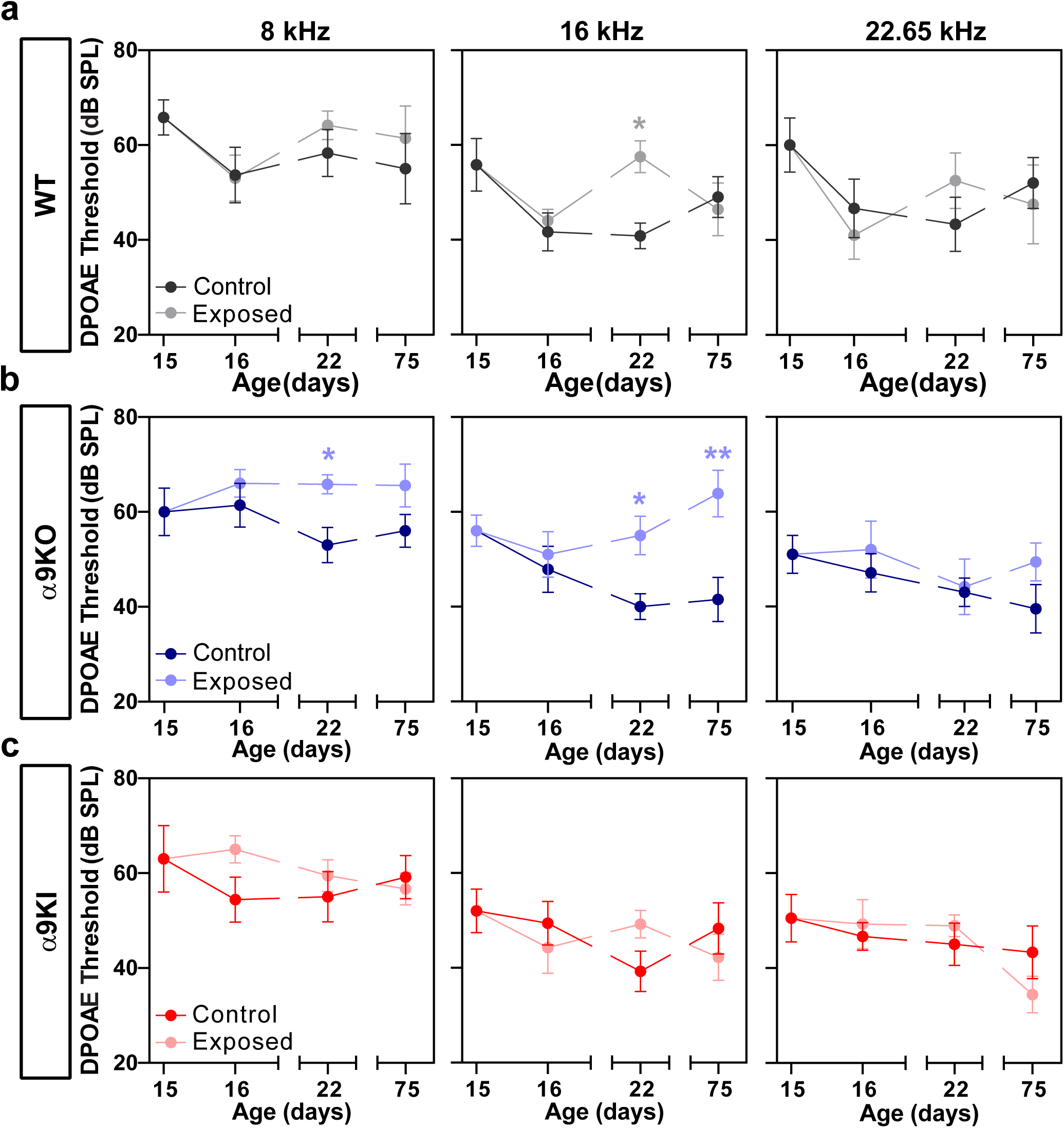
Assessment of OHC functional integrity at different postnatal ages following noise exposure at P15 in WT, α9KO, and α9KI mice. DPOAE thresholds are shown for WT (n_Control_ = 14 and n_Exposed_ = 12) (**a**), α9KO (n_Control_ = 11 and n_Exposed_ = 11) (**b**), and α9KI (n_Control_ = 12 and n_Exposed_ = 12) (**c**) at 8, 16 and 22.65 kHz. WT mice exhibited a transient increase in DPOAE thresholds 1 week post-exposure at 16 kHz. α9KO mice showed increased thresholds at 8 and 16 kHz 1 week post-exposure, with sustained elevation at 16 kHz up to 2 months. α9KI mice demonstrated no changes in DPOAE thresholds after noise exposure. Data are presented as mean ± SEM. Statistical significance is indicated by asterisks (Mann-Whitney test, *p < 0.05; **p < 0.01).

In α9KO mice, DPOAE thresholds remained unchanged one day after noise exposure. However, seven days post-exposure, significant threshold elevations were observed at 8 and 16 kHz, but not at 22.65 kHz (Mann-Whitney, df = 1, p = 0.02 at 8 kHz and p = 0.02 at 16 kHz). After two months, recovery to unexposed levels occurred only at 8 kHz, while DPOAE thresholds remained elevated at 16 kHz (Mann-Whitney, df = 1, p = 0.003 at 16 kHz; Figure 5b). In contrast, mice with enhanced MOC function showed no changes in DPOAE thresholds at any post-exposure time point (Figure 5c), suggesting that noise exposure at hearing onset did not impair the cochlear amplifier function in these mice with enhanced OHC inhibition.

### Ribbon synapse changes after early noise exposure in mice with different levels of α9α10 nAChR activity

Histological analysis was performed on cochleae dissected and fixed at P15 and P75 from unexposed mice of all three genotypes, which exhibit varying levels of α9α10 nAChR activity, to evaluate its effect of on the density and structure of IHC ribbon synapses. To visualize these ribbon synapses, we performed immunostaining on whole-mount preparations of the organ of Corti with antibodies against CtBP2-Ribeye, a presynaptic ribbon protein (Khimich et al., 2005), and GluA2, an AMPA-type glutamate receptor expressed at the postsynaptic afferent terminal (Matsubara et al., 1996; Liberman et al., 2011; Maison et al., 2013). Ribbon synapses were identified by the colocalization of CtBP2 and GluA2 puncta at the base of the IHC, as shown in representative confocal images of IHC synapses from apical whole mount organ of Corti from unexposed WT, α9KO, and α9KI mice (Figure 6a) (Liberman et al., 2011). We quantified the number of colocalized synaptic markers per IHC at both ages across three cochlear regions, each representing a distinct frequency range: apex (∼4–12 kHz), middle turn (∼12–20 kHz), and base (∼22–40 kHz) (Figure 6b). At the low-frequency apical end, both unexposed α9KO and α9KI mice showed a significant reduction in synaptic density compared to WT, with decreases of 33% and 31.18% at P15, and 30.64% and 27.94% at P75, respectively (Kruskal-Wallis, df = 2, p < 0.0001 for both groups; Figure 6b, left panel). In the middle region, synaptic density was significantly reduced in both α9KO and α9KI mice compared to WT at P15 (by 28.3% and 35.33%, respectively). However, at P75, only α9KI mice showed a significant reduction (35.97%) relative to WT (Kruskal-Wallis, df = 2, p < 0.0001; Figure 6b, middle panel). Similarly, in the high-frequency basal region, synaptic puncta were significantly reduced at P15 in both α9KO and α9KI mice (by 24.9% and 24.3%, respectively; Kruskal-Wallis, df = 2, p < 0.001 in α9KO and p < 0.01 in α9KI; Figure 6b, right panel), whereas at P75, a significant reduction of 22.07 % was observed only in α9KI mice (Kruskal-Wallis, df = 2, p = 0.02; Figure 6b, right panel). No significant changes in the number of synaptic puncta were observed between P15 and P75 in WT, α9KO, or α9KI mice across any cochlear region (Figure 6b), indicating that under unexposed conditions, alterations in transient MOC inhibition affect the number of ribbon synapses early in development and persist into maturity.

**Figure 6:**
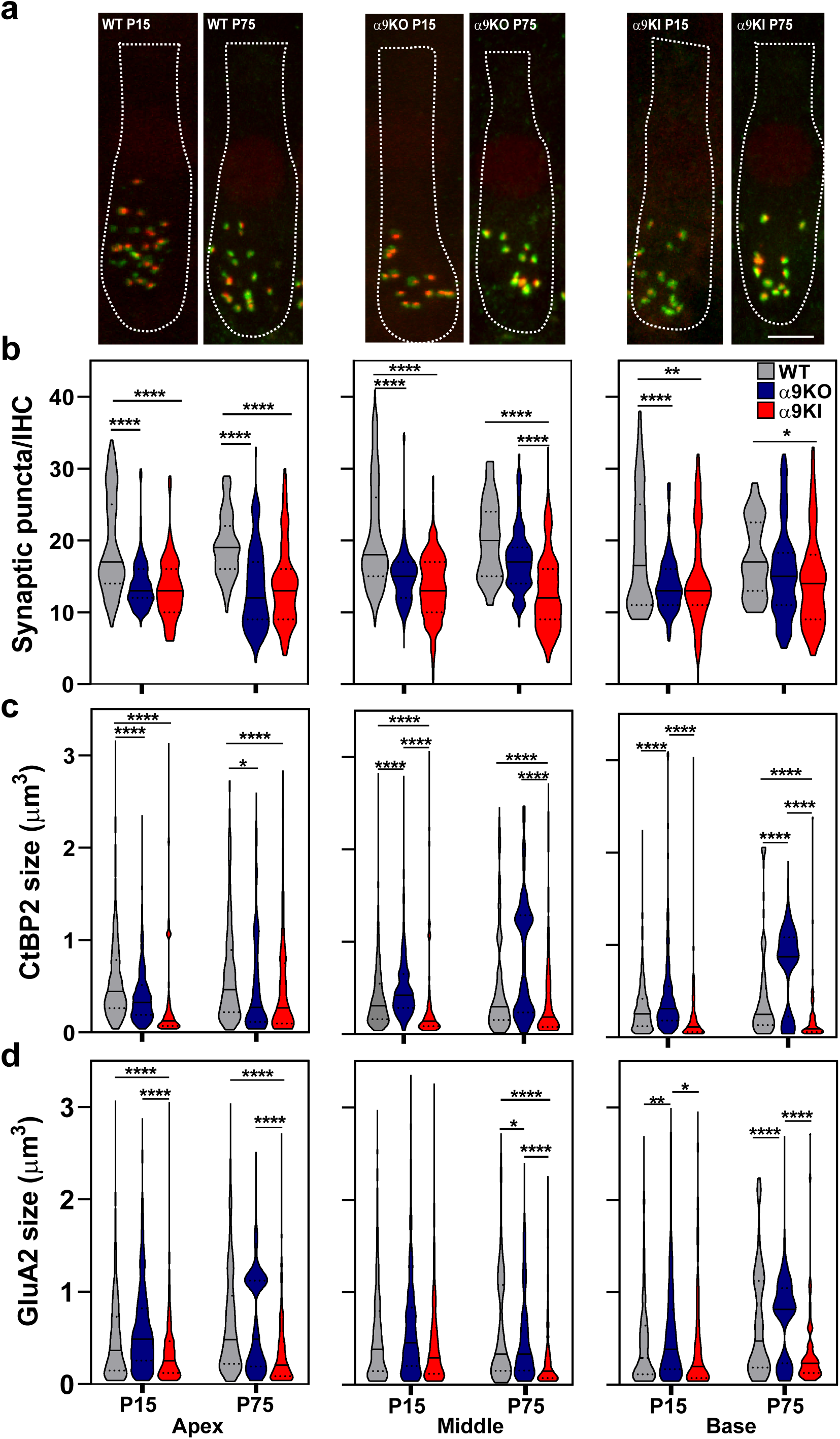
Analysis of IHC ribbon synapses in unexposed WT, α9KO, and α9KI mice at P15 and P75. (**a**) Representative confocal images of IHC synapses at the apex of the cochlea from cochleae immunolabeled for presynaptic ribbons (CtBP2-red) and postsynaptic receptor patches (GluA2-green) for WT (left), α9KO (middle) and α9KI (right) at P15 and P75. CtBP2 antibody also weakly stains IHC nuclei. Scale bar, 7 µm. (**b**) Quantification of putative ribbon synapses per IHC (i.e., colocalized CtBP2 and GluA2 puncta). WT P15: n_Apex_ = 182 IHC, n_Middle_ = 248, n_Base_ = 68 (6 animals); WT P75: n_Apex_ = 80 IHC, n_Middle_ = 75, n_Base_ = 60 IHC (14 animals); α9KO P15: n_Apex_ = 201 IHC, n_Middle_ = 230, n_Base_ = 164 (5 animals); α9KO P75: n_Apex_ = 100 IHC, n_Middle_ = 84, n_Base_ = 118 (11 animals); α9KI P15: n_Apex_ = 155 IHC, n_Middle_ = 284, n_Base_ = 85 (5 animals) and α9KI P75: n_Apex_ = 178 IHC, n_Middle_ = 319, n_Base_ = 198 (12 animals). (**c**) Volumes of fluorescence staining for ribbons (CtBP2 size) and (**d**) AMPA receptors (GluA2 size) for each of the synaptic pairs (WT P15: n_Apex_ = 684 synaptic pairs, n_Middle_ = 729, n_Base_ = 380; WT P75: n_Apex_ = 450, n_Middle_ = 509, n_Base_ = 203; α9KO P15: n_Apex_ = 845, n_Middle_ = 800, n_Base_ = 735; α9KO P75: n_Apex_ = 367, n_Middle_ = 187, n_Base_ = 221; α9KI P15: n_Apex_ = 428, n_Middle_ = 484, n_Base_ = 232 and α9KI P75: n_Apex_ = 247, n_Middle_ = 270, n_Base_ = 240). Violin plots indicate median (solid line) and interquartile range (IQR, dashed lines). Asterisks denote statistical significance. The number of synapses was analyzed using the Kruskal–Wallis test followed by Dunn’s post-test, while synaptic volumes were analyzed using one-way ANOVA followed by the Holm–Sidak test.; *p < 0.05; **p < 0.01; ***p < 0.001, ****p < 0.0001).

We further quantified the volume of CtBP2 puncta and GluA2 receptor patches within colocalized synaptic markers at both P15 and P75 across the three groups of mice (Figure 6c, d). In the apical region, both α9KO and α9KI mice showed reduced ribbon volumes compared to WT at both ages (one way ANOVA followed by Holm-Sidak test, df = 2, p < 0.0001, for both groups and ages; Figure 6c, left panel). In the middle cochlear region at P15, CtBP2 puncta were enlarged in α9KO mice and reduced in α9KI mice compared to WT. By P75, only α9KI mice continued to show a significant reduction relative to WT (one way ANOVA followed by Holm-Sidak test, df = 2, p < 0.0001, for both groups and ages; Figure 6c, middle panel). In the basal region, CtBP2 volumes were increased in α9KO mice and reduced in α9KI mice at both ages (one way ANOVA followed by Holm-Sidak test, df = 2, p < 0.0001, for both groups and ages; Figure 6c, right panel). GluA2 receptor patch size was significantly reduced in the apical region of α9KI mice compared to WT at both P15 and P75, while no changes were observed in α9KO mice (one way ANOVA followed by Holm-Sidak test, df = 2, p < 0.0001, for both ages; Figure 6d, left panel). In the middle region, no differences in GluA2 size were observed at P15; however, by P75, both α9KO and α9KI mice exhibited reduced GluA2 volumes compared to WT. (one way ANOVA followed by Holm-Sidak test, df = 2, p < 0.05 in α9KO and p < 0.0001 in α9KI; Figure 6d, middle panel). In the basal region, GluA2 puncta were significantly larger in α9KO mice compared to WT at both P15 and P75 (one-way ANOVA followed by Holm-Sidak test, df = 2, p < 0.01 at P15 and p < 0.0001 at P75; Figure 6d, right panel), while no differences were observed in α9KI mice. These results suggest that inhibitory input from cholinergic efferent fibers plays a key role in shaping synaptic density and preserving the morphology of pre-and postsynaptic elements essential for proper auditory function.

The long-term impact of noise exposure at hearing onset on auditory nerve synapse numbers is illustrated in Figure 7, on cochleae dissected and fixed at P75 from control and noise-exposed mice with varying levels of α9α10 nAChR activity. Presynaptic, postsynaptic, and colocalized synaptic puncta quantifications are shown for the apical, middle, and basal cochlear regions (Figure 7, violin plots, right panels). In WT exposed ears, the number of CtBP2, GluA2, and colocalized synaptic puncta significantly decreased at both the apical and basal cochlear ends (Figure 7a). The decrease in CtBP2 puncta was more pronounced in the high-frequency basal region, with a 37.2% decrease compared to controls (Mann-Whitney, df = 1, p < 0.0001 at the apical and basal regions). Similarly, GluA2 postsynaptic receptors were reduced in both the apical and basal turns in the cochlea of noise-exposed WT mice (Mann-Whitney, df = 1, p < 0.0001 at both cochlear regions). Putative ribbon synapse counts, defined by juxtaposed CtBP2-and GluA2-positive puncta, showed a 34.21% reduction in the apical and a 32.56% reduction in the basal turn following acoustic trauma (Mann-Whitney, df = 1, p < 0.0001 at both cochlear regions) (Figure 7a, right panel). In α9KO mice, there was also a reduction in the number of presynaptic, postsynaptic, and colocalized puncta after noise exposure, depending on cochlear frequency/location (Figure 7b). Ribbon puncta in α9KO exposed ears were reduced by up to 14.73% of the control in the apical and 24.75% in the basal cochlear ends after noise exposure (Mann-Whitney, df = 1, p = 0.001 and p = 0.002 at the apical and basal region, respectively). GluA2-positive postsynaptic receptor patches decreased by 24.79% in the basal turn of noise-exposed mice (Mann-Whitney, df = 1, p = 0.004). Following noise exposure, synaptic puncta were reduced by 26.17% in the apical cochlear region and 25.87% in the basal region (Mann-Whitney, df = 1, p = 0.002 and p < 0.0001 at the apical and basal region, respectively) (Figure 7b, right panel). Interestingly, after two months post-exposure, α9KI ears exhibited no significant changes in presynaptic ribbons, postsynaptic GluA2 receptor patches, or putative synapses across all cochlear regions (Figure 7c). Although not statistically significant, a modest increase in synaptic density (∼19% at the basal region) was observed in α9KI tissue (Figure 7c, right panel).

**Figure 7:**
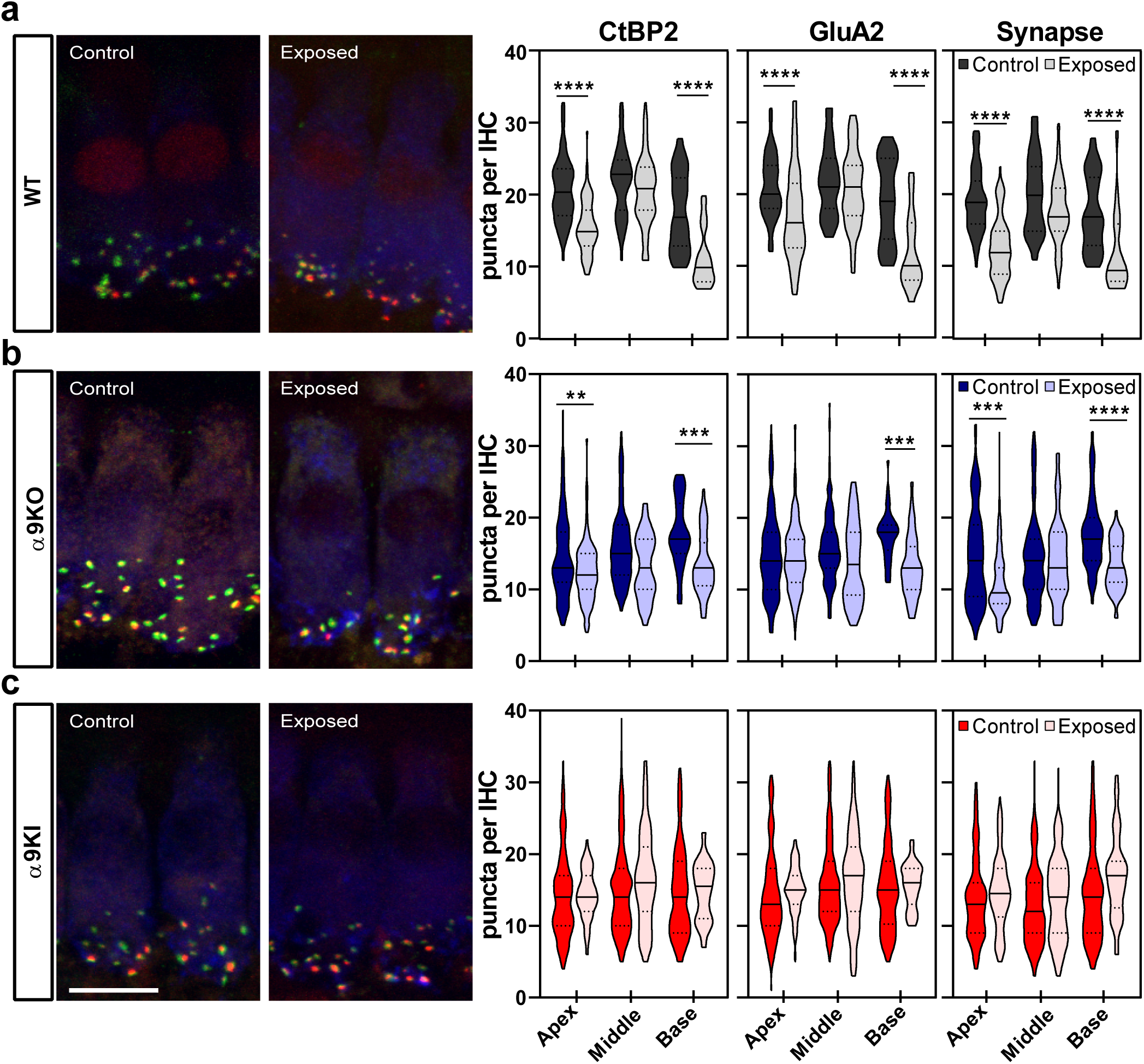
Analysis of the degree of IHC synaptopathy two months after exposure to noise at P15. Left, Representative confocal images of IHC synapses at the apex of the cochlea from cochleae immunolabeled for presynaptic ribbons (CtBP2-red), postsynaptic receptor patches (GluA2-green) and IHCs (Myosin VIIa-blue). CtBP2 antibody also weakly stains IHC nuclei. Scale bar, 10µm. Right, Quantitative data obtained from WT, α9KO, and α9KI mice at P75. For each IHC, we analyzed the number of CtBP2 puncta, postsynaptic GluA2 receptor patches, and putative ribbon synapses. (**a**) In traumatized WT mice, there was a reduction in the number of CtBP2 puncta, GluA2 receptor patches, and putative synapse, at the apical and basal region of the cochlea (WT Control: n_Apex_ = 80 IHC, n_Middle_ = 75, n_Base_= 60; WT Exposed: n_Apex_ = 82 IHC, n_Middle_ = 90, n_Base_ = 50). (**b**) In traumatized α9KO mice, there was a reduction in the number of prelocalized, postlocalized, and colocalized puncta depending on cochlear frequency/location (α9KO Control: n_Apex_ = 100 IHC, n_Middle_ = 84, n_Base_ = 118; α9KO Exposed: n_Apex_ = 234 IHC, n_Middle_ = 60, n_Base_ = 90). (**c**) In traumatized α9KI mice, no difference was found in the number of presynaptic ribbons, postsynaptic AMPA receptors, and colocalization puncta for the three regions of the cochlea (α9KI Control: n_Apex_ = 178 IHC, n_Middle_ = 319, n_Base_ = 198; α9KI Exposed: n_Apex_ = 196 IHC, n_Middle_ = 145, n_Base_ = 80). Violin plots indicate median (solid line) and IQR (dashed lines). Asterisks denote the statistical significance (Mann Whitney test, **p < 0.01; ***p < 0.001, ****p < 0.0001).

Finally, we quantified the volume of ribbons and AMPA receptor patches (i.e., colocalized CtBP2 and GluA2 puncta) in unexposed and exposed mice with different levels of MOC feedback (Figure 8). In WT mice, noise exposure resulted in significantly larger ribbon volumes in the basal end of the cochlea compared to unexposed mice, with no corresponding changes in AMPA receptor patch size (t-test, df = 1, p = 0.01; Figure 8a). No changes were observed in the apical or middle cochlear regions (Figure 8a). Conversely, α9KO mice showed reduced volumes of both ribbons and AMPA receptor patches in the basal region following noise exposure, with no changes in the apical or middle regions (t-test, df = 1, p = 0.002 for CtbP2 volume and p = 0.003 for GluA2 volume; Figure 8b). Notably, α9KI mice showed no morphological changes, suggesting that enhanced α9α10 nAChR activity protects synapses from noise-induced alterations (Figure 8c).

**Figure 8:**
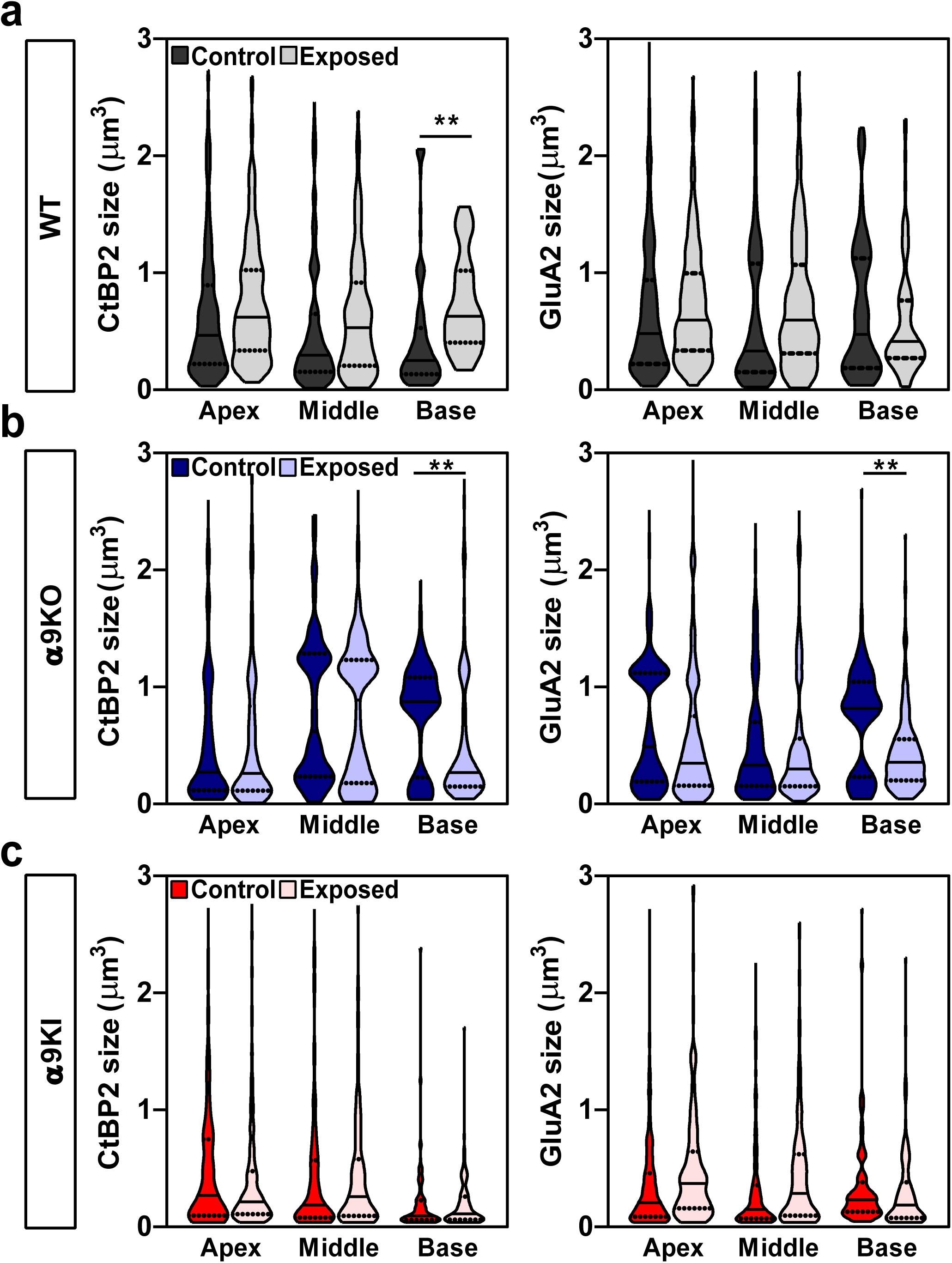
Volumes of presynaptic ribbons and postsynaptic receptor patches for each synaptic pair from control and noise-exposed WT, α9KO and α9KI mice. Violin plots comparing the volumes of fluorescence staining for ribbons (CtBP2 size) and AMPA receptors (GluA2 size) between control and noise-exposed mice. The quantification was made in the apex, middle and base regions of the cochlea from WT (**a**) (Control: n_Apex_ = 450 synaptic pairs, n_Middle_ = 509, n_Base_ = 203; Exposed: n_Apex_ = 497 synaptic pairs, n_Middle_ = 175, n_Base_ = 250); α9KO (**b**) (Control: n_Apex_ = 367 synaptic pairs, n_Middle_ = 187, n_Base_ = 221; Exposed: n_Apex_ = 207 synaptic pairs, n_Middle_ = 300, n_Base_ = 145) and α9KI (**c**) (Control: n_Apex_ = 247 synaptic pairs, n_Middle_ = 270, n_Base_ = 240; Exposed: n_Apex_ = 280 synaptic pairs, n_Middle_ = 310, n_Base_ = 360) mice. Violin plots indicate median (solid line) and IQR (dashed lines). Asterisks denote the statistical significance (t-test; *p < 0.05; ***p < 0.001).

## Discussion

Auditory development is a gradual process that begins during embryonic stages and continues through sensitive postnatal periods. While the detrimental effects of noise exposure in adulthood are well documented, less is understood about its impact during development and the role of the MOC system in this context. Here, we show that varying levels of MOC inhibition impact the timing of hearing onset and are crucial for proper development of IHC ribbon synapses. Early noise exposure caused lasting auditory deficits in WT, highlighting the greater developmental vulnerability. In contrast, enhanced cholinergic activity prevented damage, underscoring the MOC’s protective role in early development.

Before hearing begins, MOC synapses modulate IHCs’ activity, shaping the developmental increase in Ca^2+^ sensitivity of glutamate release at IHC ribbon synapses (Glowatzki and Fuchs, 2000; Johnson et al., 2011, 2013). Disruption of this input, via genetic modification or neonatal olivocochlear bundle transection, impairs cochlear and auditory circuit maturation (Walsh et al., 1998; Frank et al., 2009; Hirtz et al., 2011; Johnson et al., 2013; Clause et al., 2014; Di Guilmi et al., 2019; Babola et al., 2021). Our findings demonstrate that α9α10 nAChR activity regulates hearing onset and synaptic integrity. Enhanced α9α10 nAChR activity in α9KI mice accelerates the onset of hearing, with sound-evoked potentials detected earlier than in WT mice. In α9KO mice, which lack MOC input, hearing onset was delayed, with elevated ABR thresholds and diminished wave 1 amplitudes. This aligns with prior reports using Ca²⁺ imaging showing diminished sensitivity in α9KO inferior colliculus at hearing onset (Wang et al., 2021). Importantly, both enhanced and absent MOC inhibition disrupted IHC ribbon synapse development, with reduced synapse density and altered volumes at P15 and P75. Despite similar synaptic reductions, their distinct ABR phenotypes suggest differential effects on auditory nerve fiber subtypes. The earlier hearing onset in α9KI mice could reflect preservation or accelerated maturation of low-threshold, high-spontaneous-rate fibers, while their disruption in α9KO mice may contribute to delayed onset. Morphologically, α9KI mice showed reduced ribbon and GluA2 patch volumes across all cochlear turns, whereas α9KO mice exhibited region-specific changes: smaller ribbons in apical regions with unchanged GluA2 patches, and enlarged pre-and post-synaptic components in middle/basal turns. These ribbon alterations may reflect disruption of Ca²⁺-dependent ribbon fusion or stabilization, impairing synaptic refinement. Ribbon size and maturation is tightly regulated by presynaptic Ca^2+^ influx (Frank et al., 2009; Sheets et al., 2012; Wong et al., 2019; Özçete and Moser, 2021). Fusion of ribbon precursors during this period is thought to support sustained release at mature active zones (Michanski et al., 2019). In developing IHCs, intracellular Ca^2+^ dynamics are shaped by voltage-gated Ca²⁺ channels and by α9α10 nAChRs, which are highly permeable to Ca²⁺ (Weisstaub et al., 2002; Gomez-Casati et al., 2005; Moglie et al., 2018). In α9KO mice, the absence of α9α10-mediated Ca²⁺ influx leads to increased action potential frequency and disrupted exocytosis (Johnson et al., 2013). In α9KI mice, the L9’T mutation reduces receptor desensitization, likely prolonging Ca²⁺ entry and altering IHC firing during the prehearing period (Taranda et al., 2009). These results highlight the MOC synapse’s role in coordinating afferent and efferent activity for normal auditory development.

Auditory sensitivity in vertebrates matures from high thresholds and poor tuning at hearing onset to precise, low-threshold processing (Song et al., 2006). In WT mice, ABR and DPOAE data from P12 to P75 showed this progression. At P14, thresholds were elevated and waveforms immature, but by P22, thresholds and wave 1 amplitudes reached mature levels. Thus, at P15, the time of noise exposure, auditory maturation was still incomplete. A key finding of this study is that noise exposure during early development leads to more severe and lasting effects than those reported at later stages. In WT mice, noise exposure at P15 caused lasting threshold shifts and 35% synapse loss, while the same exposure at P21 led to only transient threshold shifts and milder synaptopathy (Boero et al., 2018). This underscores the heightened vulnerability at hearing onset, likely due to both noise-induced damage and disruption of ongoing synaptic or hair cell maturation. Similar age-dependent vulnerability has been reported in other juvenile models (Fernandez et al., 2015; Jensen et al., 2015; Schrode et al., 2022; Lu et al., 2024). While the mechanisms behind this heightened sensitivity remain unclear, it may arise from ongoing maturation of the MOC system and central auditory pathways, including myelination and afferent input. Developing interactions between the auditory nerve and LOC system may further increase susceptibility to noise exposure.

In WT mice, DPOAE thresholds were elevated seven days post-exposure, reaching statistical significance at 16 kHz, but returned to baseline by two months, suggesting preserved or repaired OHC function. Whether this reflects true recovery, delayed maturation, or compensation remains to be determined. In contrast, persistent ABR threshold elevation and reduced wave 1 amplitudes suggest lasting afferent damage, likely due to cochlear synaptopathy affecting both high-and low-threshold auditory nerve fibers (Reijntjes et al., 2025). This is the first report of sustained ABR threshold elevation following moderate noise exposure, similar to lasting high-frequency shifts and presynaptic loss seen in P14 rats and noise-sensitive P15 C57BL/6J mice (Davis et al., 2003) exposed to high-intensity sound (Rybalko et al., 2015; Boero et al., 2021). Noise exposure caused significant ribbon synapse loss in apical and basal cochlear regions of WT mice, correlating with reduced ABR wave 1 amplitudes; however, despite functional deficits at 16 kHz, synaptic loss in the mid-frequency region was minimal—reflecting a known mismatch due to cochlear place-frequency map shifts after noise exposure (Müller and Smolders, 2005). In addition, IHCs of noise-exposed mice tend to have larger ribbons, especially in the basal turn. Similar post-trauma ribbon enlargement was observed in RIBEYE-tagRFP mice (Mohamad et al., 2024) and via volume EM (Lu et al., 2024), likely reflecting the fusion of individual ribbons. Multi-ribbon active zones and ribbon unanchoring from synaptic tethers have been observed in synaptopathic ears (Moverman et al., 2023; Lu et al., 2024; Mohamad et al., 2024). Though their function is unclear, these structures persist into adulthood (Michanski et al., 2019; Hua et al., 2021) and are a hallmark of cochlear aging (Stamataki et al., 2006), with larger ribbons being associated with increased exocytotic activity (Jeng et al., 2020). Notably, noise exposure has also been associated with enhanced exocytosis in IHCs, along with an increase in tone-burst responses in auditory nerve fibers (Boero et al., 2021; Suthakar and Liberman, 2021).

In α9KO mice, ABR and DPOAE thresholds were elevated one week post-exposure at P15, consistent with prior findings at P21 (Boero et al., 2018). However, unlike WT, ABR thresholds in α9KOs nearly fully recovered by two months. DPOAE thresholds also normalized except at 16 kHz, matching a slight, non-significant ABR threshold elevation at that frequency. The recovery of ABR thresholds observed in α9KO mice, unlike the persistent elevation in WT mice, may stem from reduced auditory sensitivity at hearing onset in α9KO mice, leading to less effective noise exposure and milder impact. Despite threshold recovery, reduced wave 1 amplitudes and synapse counts suggest lasting damage to high-threshold, low-spontaneous-rate fibers (Furman et al., 2013; Reijntjes et al., 2025). Additionally, noise-exposed α9KO ears showed reduced sizes of pre-and postsynaptic puncta, indicating structural synaptic alterations. At P16 (one day post-exposure), ABR and DPOAE thresholds in both WT and α9KO mice were slightly reduced compared to P15 and aligned with those of unexposed controls, suggesting that noise had no immediate impact on auditory maturation at this stage. In contrast, Boero et al. (2021) reported lasting damage in C57BL/6J mice one day after 120 dB SPL at P15, highlighting that early noise effects depend on complex interactions between factors such as intensity, duration, and genetic background, and differ from outcomes seen with noise exposure at mature stages.

Unlike WT and α9KO mice, α9KI mice showed no threshold shifts or synaptopathy after noise, consistent with prior reports in mature α9KI mice (Taranda et al., 2009; Boero et al., 2018) and recent gene therapy studies supporting protective effects of enhanced α9α10 nAChR activity (Zhang et al., 2023; Slika et al., 2025). The protective role of the MOC system has been recognized for decades: efferent stimulation reduces threshold shifts (Rajan, 1988; Reiter and Liberman, 1995), while surgical de-efferentation increases vulnerability (Kujawa and Liberman, 1997). Overexpressing wild-type α9 also confers greater resistance to noise (Maison et al., 2002). *In vivo* studies show that ACh evokes fast (∼100ms) and slow (∼10s) MOC effects on cochlear responses (Sridhar et al., 1995, 1997; Cooper and Guinan, 2003). Fast suppression involves SK2-mediated OHC hyperpolarization, while slow suppression likely reflects Ca^2+^ signaling and increased OHC stiffness, both reducing cochlear vibrations (Cooper and Guinan, 2003). Evidence suggests that efferent protection likely relies on slow, Ca²⁺-dependent mechanisms that reduce acoustic injury by dampening cochlear vibrations or altering OHC function via α9α10 receptor-mediated Ca²⁺ influx (Reiter and Liberman, 1995; Maison et al., 2007). Though MOC-OHC innervation is still maturing during early development (Simmons et al., 1996), our findings show that enhanced α9α10 activity already confers robust protection against acoustic trauma.

